# A full spectrum PNGase activity analysis of R328 mutations on NGLY1

**DOI:** 10.1101/2022.04.07.487431

**Authors:** Shuying Yuan, Yanwen Chen, Lin Zou, Xinrong Lu, Ruijie Liu, Shaoxing Zhang, Li Chen, Guiqin Sun

**Affiliations:** School of Medical Technology and Information Engineering, Zhejiang Chinese Medical University, Hangzhou, Zhejiang 310053, China; Department of Medical Microbiology and Parasitology, Key Laboratory of Medical Molecular Virology of Ministries of Education and Health, School of Basic Medical Sciences, Fudan University, Shanghai 200032, China

**Keywords:** NGLY1-CDDG, PNGase, R328, mutation, deglycosylation, enzymatic activity

## Abstract

In humans, N-glycanase 1 (NGLY1; Peptide: N-glycanase, PNGase) is responsible for the deglycosylation of misfolded glycoproteins. Pathogenic mutations in *NGLY1* cause a clinical condition known as congenital disorder of deglycosylation (NGLY1-CDDG), a rare autosomal recessive disease first reported in 2012. Although NGLY1-CDDG was diagnosed through whole-exome or whole-genome sequencing and by evaluating the expression levels of NGLY1, the clinical relevance of a detected mutation in *NGLY1* needs to be further confirmed. In this study, an in vitro enzymatic assay system was established to evaluate the thermal stability and substrate specificity of NGLY1, as well as the optimum reaction conditions for its activity. A panel of all mutations at the amino acid site R328 in NGLY1 was subjected to this assay. The results revealed that R328A, R328D, R328E, R328F, R328G, R328I, R328P, R328V, R328W, and R328Y were dysfunctional mutations (10/19); NGLY1 mutations with R328H and R328T exhibited similar activity as wild-type NGLY1 (2/19); and NGLY1 mutations with R328C, R328K, R328L, R328M, R328N, R328Q, and R328S showed decreased activity (7/19) compared to wild-type NGLY1. In addition, the effect of potential regulatory compounds, including N-acetyl-L-cysteine and dithiothreitol, on NGLY1 was examined. This in vitro assay may serve as a standard protocol to facilitate rapid diagnosis of all mutations on NGLY1 and a practical screening method for drugs and compounds with potential therapeutic value for NGLY1-CDDG patients.

## Introduction

The human cytosolic peptide:N-glycanase (PNGase; N-glycanase 1, NGLY1), encoded by *NGLY1* gene (OMIM* 610661), acts on N-glycoproteins to generate free N-glycans and deglycosylated peptides (Suzuki et al., 2016). *NGLY1* mutation causes a congenital disorder of deglycosylation (NGLY1-CDDG), a rare human autosomal recessive disease (Enns et al., 2014; Need et al., 2012). Almost 80 NGLY1-CDDG patients were found worldwide, including 6 cases reported in China (Abuduxikuer et al., 2020; Haijes et al., 2019) (NGLY1 deficiency handbook, www.NGLY1.org). NGLY1-CDDG is a multisystemic neurodevelopmental disorder in which individuals exhibit typical features, such as developmental delay, multifocal epilepsy, microcephaly, intellectual disability, liver dysfunction, alacrima, hypohidrosis, and movement disorder (Abuduxikuer et al., 2020; Lipinski et al., 2020). However, identifying NGLY1-CDDG is a challenge for clinicians owing to the lack of a specific clinical phenotype (Haijes et al., 2019). Two potential biomarkers of NGLY1-CDDG have been described, including the N-glycoprotein structure of Neu5Ac1Hex1GlcNAc1-Asn detected in urine reported by Haijes et al. (Hall et al., 2018) and the increased aspartyl-glycosamine content in blood reported by Hall et al. (Haijes et al., 2019). Previously, NGLY1-CDDG patients were diagnosed based on whole-exome sequencing (WES) or whole-genome sequencing (WGS) data or the levels of NGLY1 in serum(Abuduxikuer et al., 2020; Need et al., 2012; Rios-Flores et al., 2020).

NGLY1-CDDG patients harbor various *NGLY1* mutations. Mutations in the *NGLY1* gene might affect the biological function of NGLY1 in vivo, leading to disease. Our group reported, for the first time, a mutation in NGLY1 wherein arginine-328 (R328) was mutated to glycine (R328G), which was a novel functional, active site in NGLY1 (Abuduxikuer et al., 2020). We also found an arginine to leucine mutation at 328 site (R328L) in another NGLY1-CDDG patient (unpublished). Recently, Dabaj et al. reported an NGLY1-CDDG patient harboring an arginine to cysteine mutation (R328C) (Dabaj et al., 2021). We analyzed the amino acid sequence of NGLY1 from *Homo sapiens, Pan troglodytes, Mus musculus, Danio rerio, Gallus gallus, Drosophila melanogaster, Caenorhabditis elegans, Saccharomyces cerevisiae*, and *Schizosaccharomyces pombe* and showed that the NGLY1-R328 site was a conserved site. Furthermore, Katiyar et al. reported that the R210 site (corresponding to position R328 of NGLY1) of the yeast peptide:*N-glycanase* (Png1p; PNGase) was important for folding and structural stability of the enzyme (Katiyar et al., 2002). Therefore, we speculated that the NGLY1-R328 site mutating to other amino acids might be a potential pathogenic mutation leading to NGLY1-CDDG. However, it was unclear whether these mutations would cause the dysfunction in NGLY1; thus, a relationship between genotype (*NGLY1* mutation) and phenotype (NGLY1 activity) needs to be established.

In this study, an in vitro standard assay for determining enzymatic activity was established. Considering R328 as an example, full-spectrum amino acid mutation libraries of the R328 site to the other 19 amino acids were constructed. Based on the in vitro enzymatic assay, purified 19 NGLY1 mutations were tested. A relationship between genotype and phenotype in NGLY1-R328 was established. In addition, the effects of chemicals and metal ions on NGLY1 were analyzed. This in vitro assay might provide a quick diagnosis for all mutations in NGLY1 and a practical screening method for drugs and compounds with potential therapeutic value for NGLY1-CDDG patients.

## Methods

### Prokaryotic expression and purification of NGLY1

To determine NGLY1 activity in vitro, the NGLY1 protein was expressed and purified using a standard method (Abuduxikuer et al., 2020; Sun et al., 2015). *NGLY1* cDNA was cloned into the pET-28a (+) plasmid between the NcoI and XhoI cloning sites. The recombinant plasmid was verified by sequencing for prokaryotic expression and transformed into *Escherichia coli* BL21 (DE3) cells. The transformants were cultured at 37 °C for 12–16 h in Luria-Bertani (LB) medium containing 50 μg/mL kanamycin to obtain isolated colonies.

Single colonies were inoculated into 5 mL of LB broth and incubated at 37 °C at 180 rpm for 12–16 h. Thereafter, the bacterial suspension was diluted with LB broth (50 μg/mL kanamycin) at a ratio of 1:100 and cultured at 37 °C at 180 rpm (OD_600_ = 0.6). IPTG (1.0 mM) was added to induce protein expression at 28 °C, and the culture was further incubated for 12 h. The cells were harvested by centrifugation at 3 500 rpm for 10 min, washed three times with PBS, and resuspended in Lysis Buffer (20 mM sodium phosphate, 300 mM sodium chloride, 10 mM imidazole, pH 7.4). The resuspended cells were sonicated (program: 5 s of sonication followed by a 10 s pause per cycle, 120 W) for 30 min on ice. After centrifugation at 12,000 rpm for 30 min at 4 °C, the protein-containing supernatant was loaded onto HisPur™ Ni-NTA Spin Columns (Thermo Fisher Scientific, USA) and purified for 3 h at 4 °C. The expressed protein was loaded onto a Ni-NTA Spin column, washed with two resin-bed volumes of Wash Buffer (20 mM sodium phosphate, 300 mM sodium chloride, 25 mM imidazole, pH 7.4) three times, and eluted by one resin-bed volume of Elution Buffer (20 mM sodium phosphate, 300 mM sodium chloride, 250 mM imidazole, pH 7.4) three times. The concentration of the purified protein was determined using a Nanodrop One (Thermo Fisher Scientific).

### Enzymatic activity assay of NGLY1

The standard N-glycoprotein bovine pancreatic ribonuclease B (RNase B, New England Biolabs, USA) was used as a substrate to measure NGLY1 activity. Oligosaccharides of RNase B cleaved by NGLY1 were analyzed using MALDI-TOF mass spectrometry (MS) and capillary electrophoresis (CE) (Cao et al., 2020; Cong et al., 2020; Sun et al., 2015; Zhang et al., 2011). In addition, enzymatic activity was also analyzed using SDS-PAGE (Abuduxikuer et al., 2020).

For each reaction, 100 μg of RNase B in phosphate-buffered saline (PBS, 137 mM sodium chloride, 2.7 mM potassium chloride, 10 mM dibasic sodium phosphate, 2 mM potassium dihydrogen phosphate, and pH 7.4) was denatured at 100 °C for 10 min and then mixed with 100 μg of NGLY1. The reaction was conducted in a 37 °C water bath for 12 h. The reaction products were desalted using a Supelclean™ Solid Phase Extraction Tube (Sigma-Aldrich, USA). The released glycans were collected by ultrafiltration using a filter with a molecular weight cut-off of 3,000 (Millipore, USA) at 12,000 rpm for 30 min at 4 °C. Finally, the products were gathered through a vacuum dryer and dissolved in 160 µL of ddH_2_O.

For MALDI-TOF MS analysis, 1 µL of the filtered solution was deposited on a MALDI plate with 1 μL of saturated 2,5-dihydroxybenzoic acid using the dried droplet method for MS analysis. Glycan profiling was performed in the positive ion reflectron mode in a rapifleX MALDI-TOF mass spectrometer (Bruker, Germany), using a nitrogen-pulsed laser (337 nm) and an acceleration voltage of 20 kV (Sun et al., 2015; Zhang et al., 2011).

For CE analysis, the sample was derivatized with 8-amino-1,3,6-pyrenetrisulfonic acid (APTS; Molecular Probes, USA). The labeled N-glycans were analyzed through DNA sequencer-assisted, fluorophore-assisted carbohydrate electrophoresis (DSA-FACE) technology using a CE-based ABI 3500 Genetic Analyzer (Applied Biosystems, USA). Data were analyzed using the GeneMapper software version 4.1 (Cao et al., 2020).

The N-glycans of serum from healthy controls were also analyzed by the CE method, as previously reported (Cao et al., 2020; Cong et al., 2020). The N-linked glycans present on the serum proteins in 2 mL of healthy serum were released by PNGase F and NGLY1, respectively, and detected using the same protocol described above. In addition, serum glycoprotein N-glycan profiles were obtained following the instructions of the Glycan-Test Kit (Sysdiagno Biomedtech, China).

### Enzymatic reaction conditions for NGLY1

The enzyme digestion reaction was performed in a 20 μL mixture containing 2 μg of denatured substrates, 2 μg of NGLY1, and PBS added up to 20 μL at 37 °C in a water bath for 12 h. The following factors were evaluated in the abovementioned reaction conditions:

#### Substrate specificity

The following specific substrates were used: RNase B with N-linked high-mannose oligosaccharide, ovalbumin (OVA) with N-linked hybrid oligosaccharide (LABELED, China), and self-made immunoglobulin G (IgG) with N-linked complex oligosaccharide.

#### Effect of temperature

NGLY1 and RNase B were mixed and incubated at 4 °C, 25 °C, 35 °C, 45 °C, 55 °C, 65 °C, 75 °C, and 85 °C.

#### Effect of pH

NGLY1 and RNase B were mixed and then added to buffer at varying pH values from 3.5–9.0 (0.1 mM citric acid and trisodium citrate, pH = 3.5–5.5; 0.1 mM PBS, pH = 6.5–8.5; 0.05 mM borax and sodium borate, pH = 9.0).

#### Thermal stability

NGLY1 was preincubated at 4 °C, 25 °C, 35 °C, 45 °C, 55 °C, 65 °C, 75 °C, and 85 °C for 1 h prior to the addition of RNase B.

#### Effect of chemical compounds

The reaction mixture was supplemented with additional chemical compounds in a 37 °C water bath for 12 h. N-acetyl-L-cysteine (NAC: 0.1, 0.5, 1.0, and 5 mM), dithiothreitol (DTT: 5, 10, 20, 30, 40, and 50 mM), sodium dodecyl sulfate (SDS, % and 0.5%), benzyloxycarbonyl-Val-Ala-Asp-fluoromethyl ketone (Z-VAD-FMK, 0.1 and 0.5 mM) and hypochlorous acid (0.001% and 0.005%).

#### Effect of mental ions

MgCl_2_, CaCl_2_, BaCl_2_, and ZnCl_2_ were added at concentrations ranging from 0.1 to 1 mM to examine their effects on NGLY1 activity.

Relatively quantitative analysis of enzyme activity-Enzyme: The relative enzymatic activity was assessed using the ImageJ software.

### Construction of full-spectrum amino acid mutation libraries on R328 site mutations of NGLY1

Primers for 19 mutations were designed using QuikChange Primer Design (https://www.agilent.com/store/primerDesignProgram.jsp)) (Table 1). The mutations were obtained using a PCR-based mutagenesis protocol following previously reported studies (Abuduxikuer et al., 2020). Each 50 µL reaction consisted of 25 µL of PrimeSTAR® Max DNA Polymerase (Takara, Japan), 21 µL of RNase-free water, 1.5 µL of each primer (forward and reverse, both at 10 µM), and 1 µL of recombined plasmid *NGLY1*/*pET28a*. Cycling times and temperatures were 98 °C for 3 min, followed by 24 cycles of 98 °C for 30 s, 60 °C for 1 min, and 68 °C for 10 s (thermal cycling). All the mutated plasmids were confirmed by sequencing and then transformed into *E. coli* BL21 (DE3) cells.

**Table 1.**
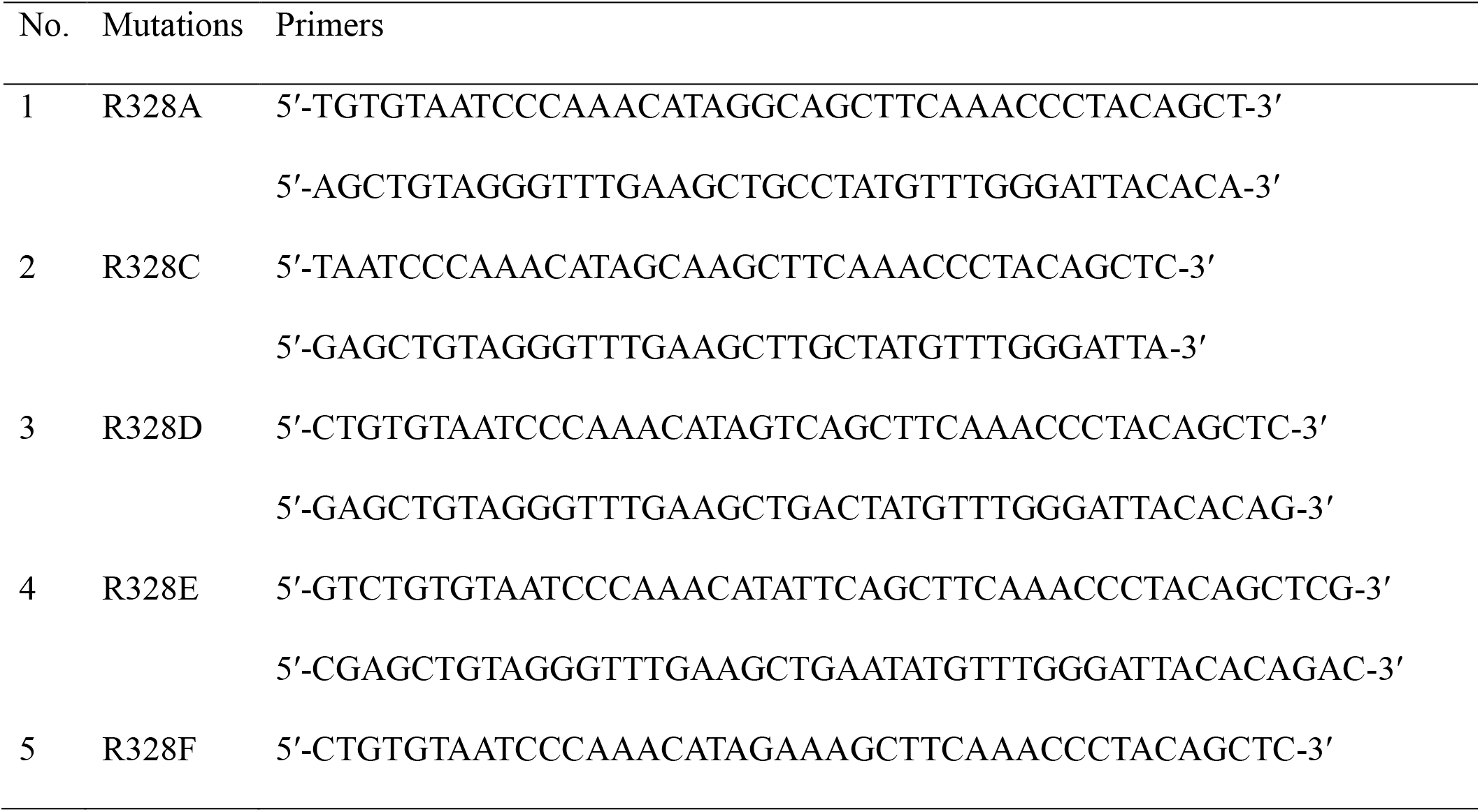
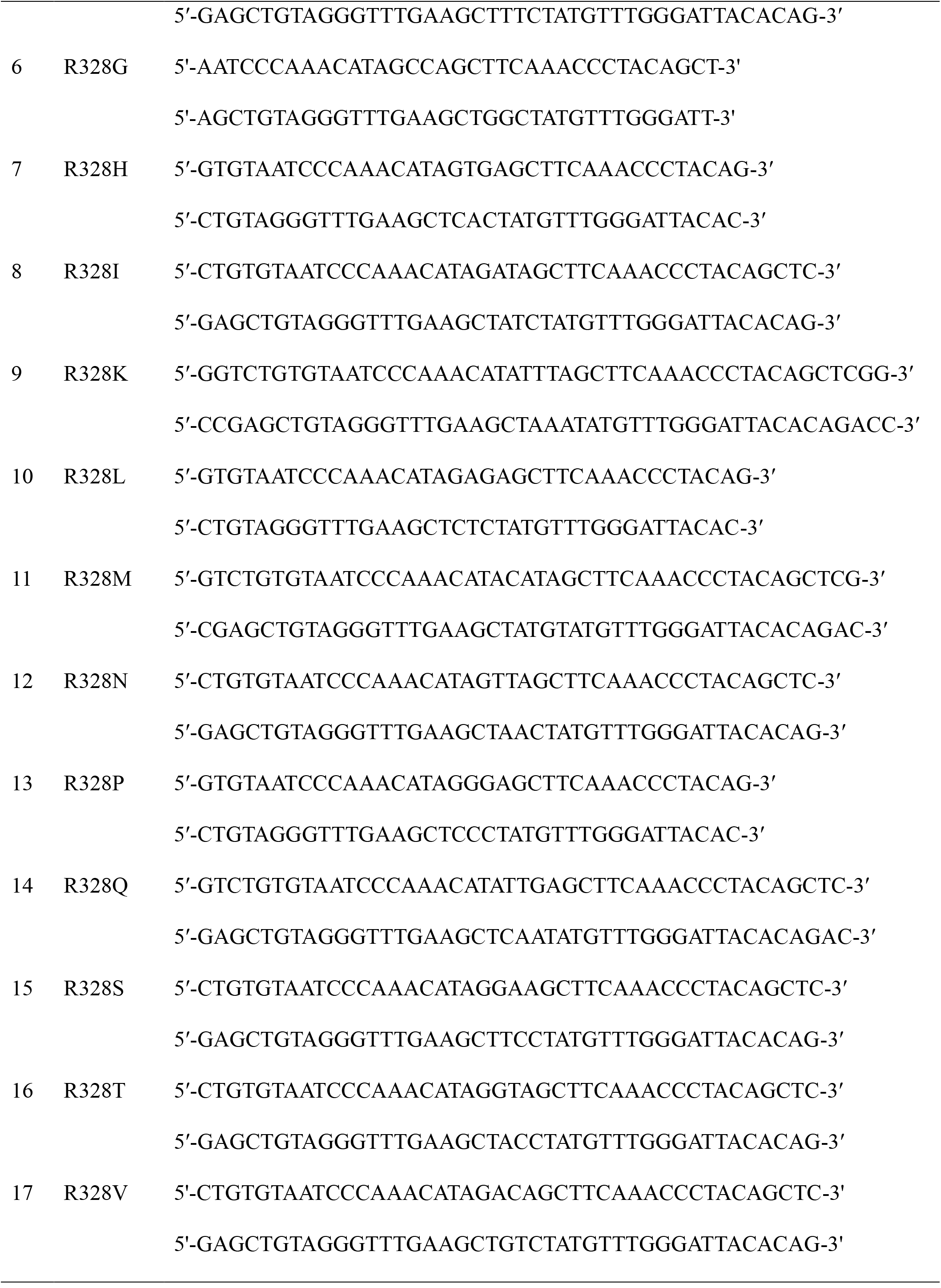
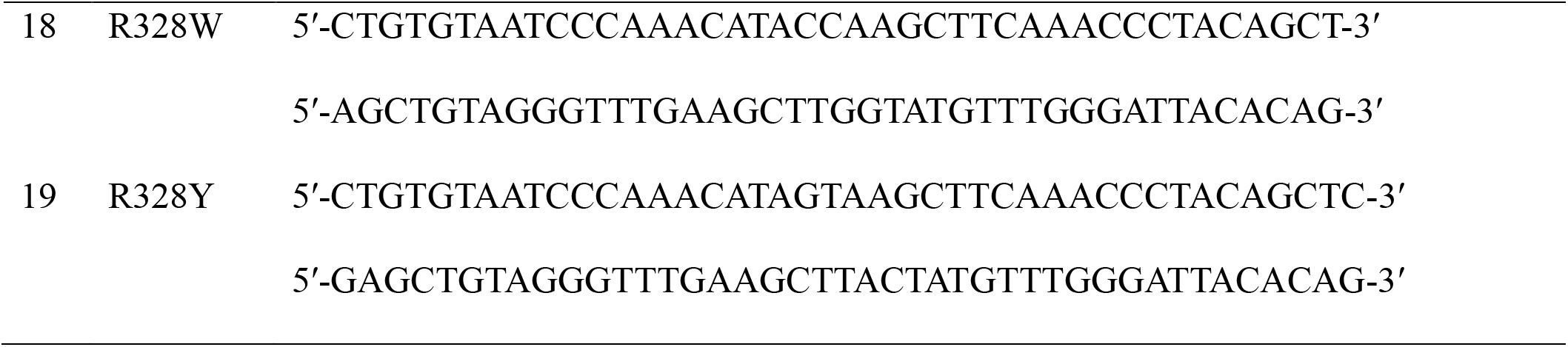
Primers of R328 NGLY1 mutations

### Enzymatic activity assay of 19 mutations in the NGLY1-R328 site

RNase B was chosen as a substrate for measuring the activity of mutated NGLY1 and analyzed by SDS-PAGE. The reaction condition for qualitative enzymatic activity of NGLY1-R328 mutations was referred to the wild type NGLY1. For those remained-active mutations, 1 μg RNase B and 2 μg of mutated NGLY1 were mixed in 20 μL reaction system. The reaction was conducted at 37 °C in a water bath for 1 h to determine the relative enzyme activity.

### Simulated structure of NGLY1

NGLY1 comprises three domains: PNGase- and ubiquitin-related (N-terminal, PUB domain), PNG Core, and PNGase and other worm proteins (C-terminal, PAW domain). The PUB domain is associated with the ubiquitin-proteasome system, PAW is involved in the oligosaccharide binding of PNGase, and the PNG Core domain is the core catalytic domain (Suzuki, 2015). The protein structure prediction of NGLY1 and 19 mutations of NGLY1 was performed using AlphaFold2 (Jumper et al., 2021; Mirdita et al., 2021). In addition, the amino acid side chains of NGLY1 and its mutations were analyzed using the PyMOL software.

## Results

### Enzymatic activity of prokaryotic NGLY1

The enzymatic activity of purified NGLY1 was evaluated using RNase B as a substrate. PNGase F was used as a positive control. The profile of the released glycans from RNase B treated with NGLY1 was detected through MALDI-TOF MS (Fig. 1A), consistent with the reported N-glycan structures (Sun et al., 2015; Wang et al., 2018b). The results of CE also showed that the profiles of glycans released from RNase B were the same as those after treatment with NGLY1 or PNGase F (Fig. 1B). RNase B digestion was verified by the downward shift of the RNase B glycoprotein band observed during SDS-PAGE analysis (upper right picture of Fig. 1A). The results indicated that the prokaryotic NGLY1 possessed similar activity as PNGase F: both hydrolyzed the intact glycans from RNase B.

**Figure 1.**
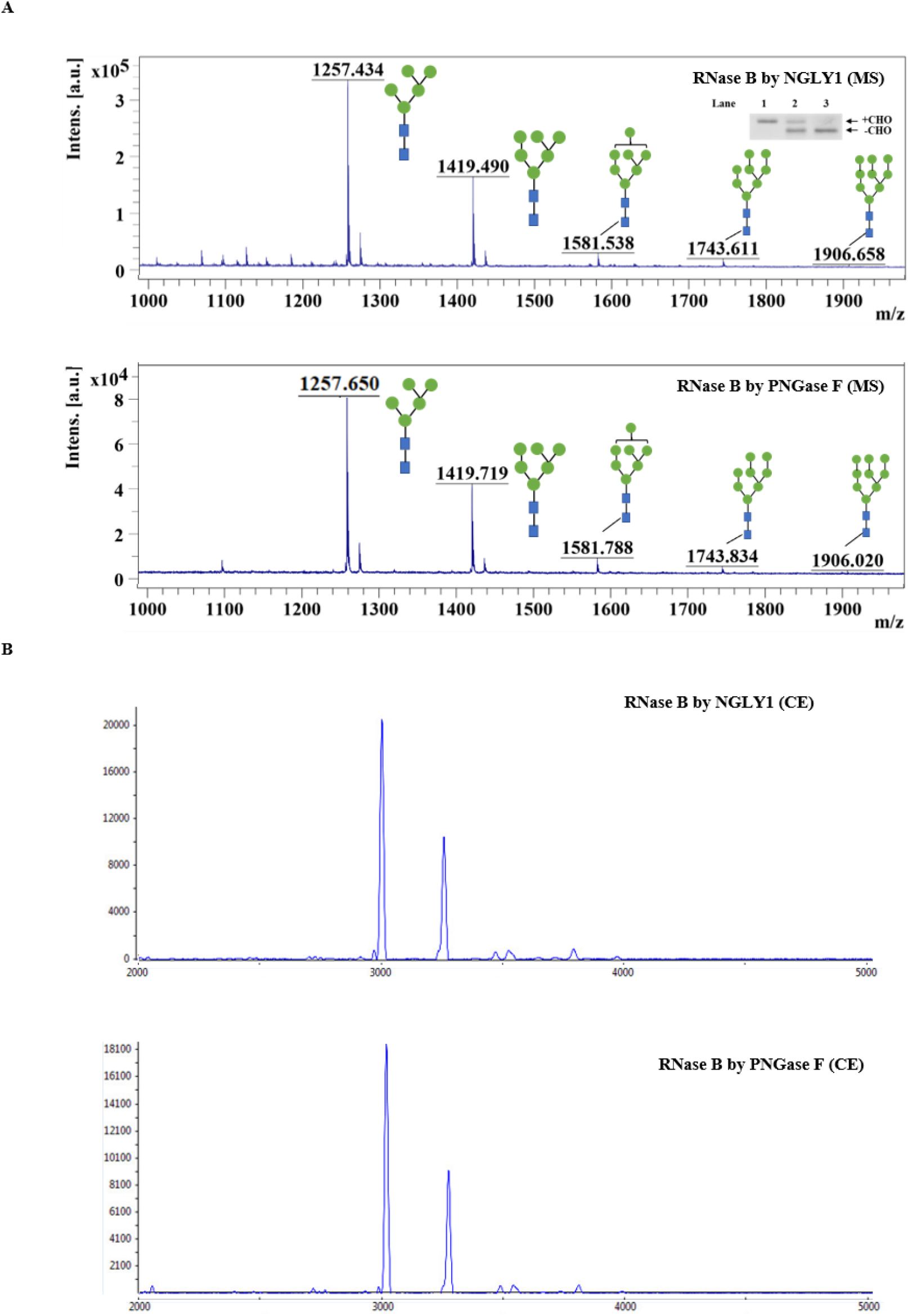
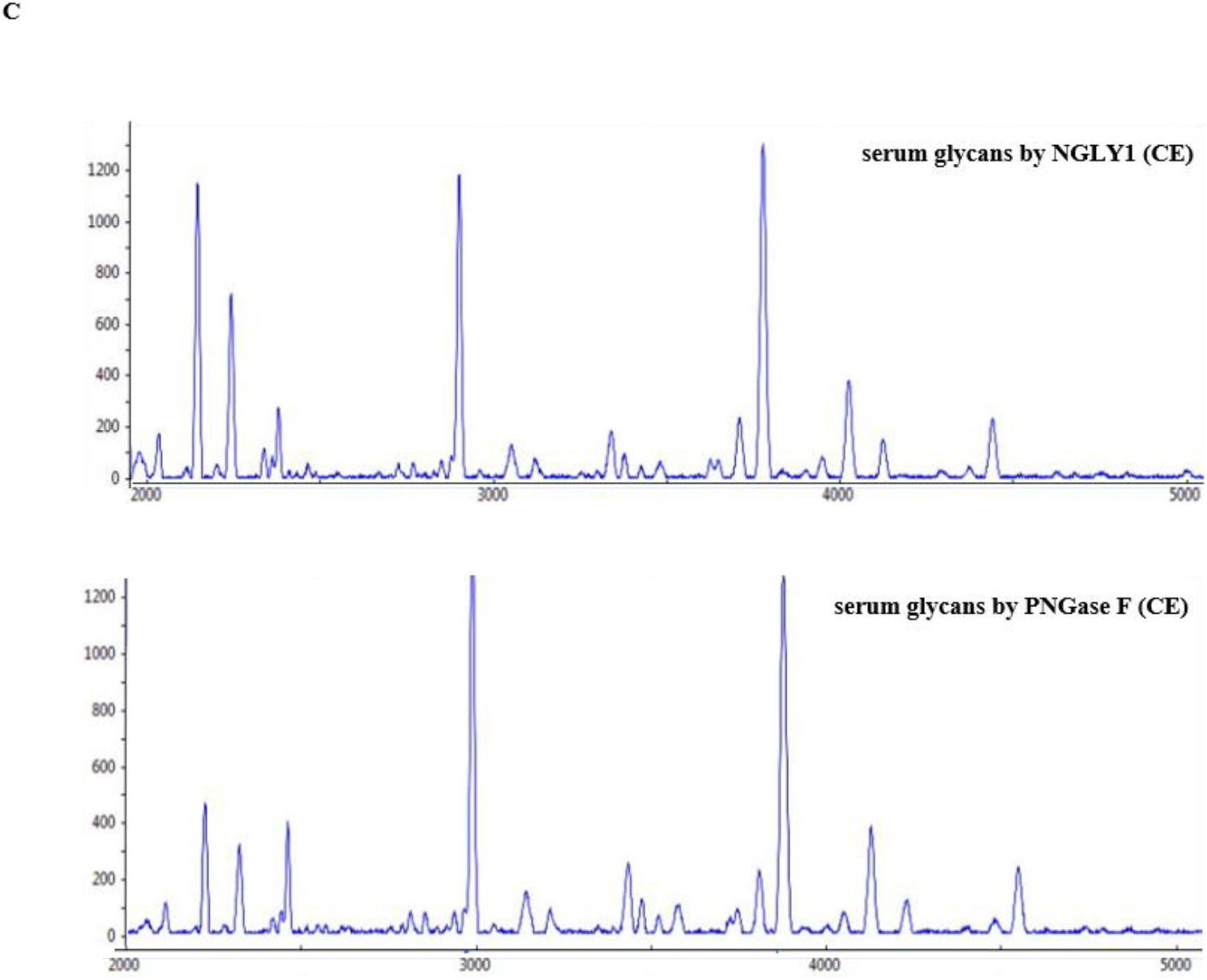
The glycosidase activity analysis of NGLY1. (A) The released N-glycans from RNase B were detected by MALDI-TOF MS. Upper right image: SDS-PAGE analysis of RNase B treated with NGLY1. Comparison of the deglycosylation activities of NGLY1 and PNGase F (glycosylated and deglycosylated forms of the substrates are indicated by +CHO and -CHO, respectively). Lane 1: RNase B, Lane 2: NGLY1 + RNase B; Lane 3: PNGase F + RNase B. (B) The released N-glycans from RNase B detected by CE. (C) The released N-glycans from total serum glycoproteins of healthy individuals detected by CE.

The function of NGLY1 has been proven by the release of N-glycans on healthy serum glycoproteins through the CE method (Fig. 1C). In addition, the N-glycans from serum glycoproteins were similarly treated with NGLY1 and PNGase F. The results indicated that NGLY1 could be used as an alternative tool for glycomics.

### Establishing the in vitro NGLY1 enzymatic assay system

The effects of temperature and pH were verified to optimize the in vitro enzymatic assay (Fig. 2). The results showed that the enzymatic activity was maintained at the reaction temperature range of 10 °C to 45 °C (Fig. 2B). NGLY1 was active at pH between 6.5 and 8.0 (Fig. 2C).

**Figure 2.**
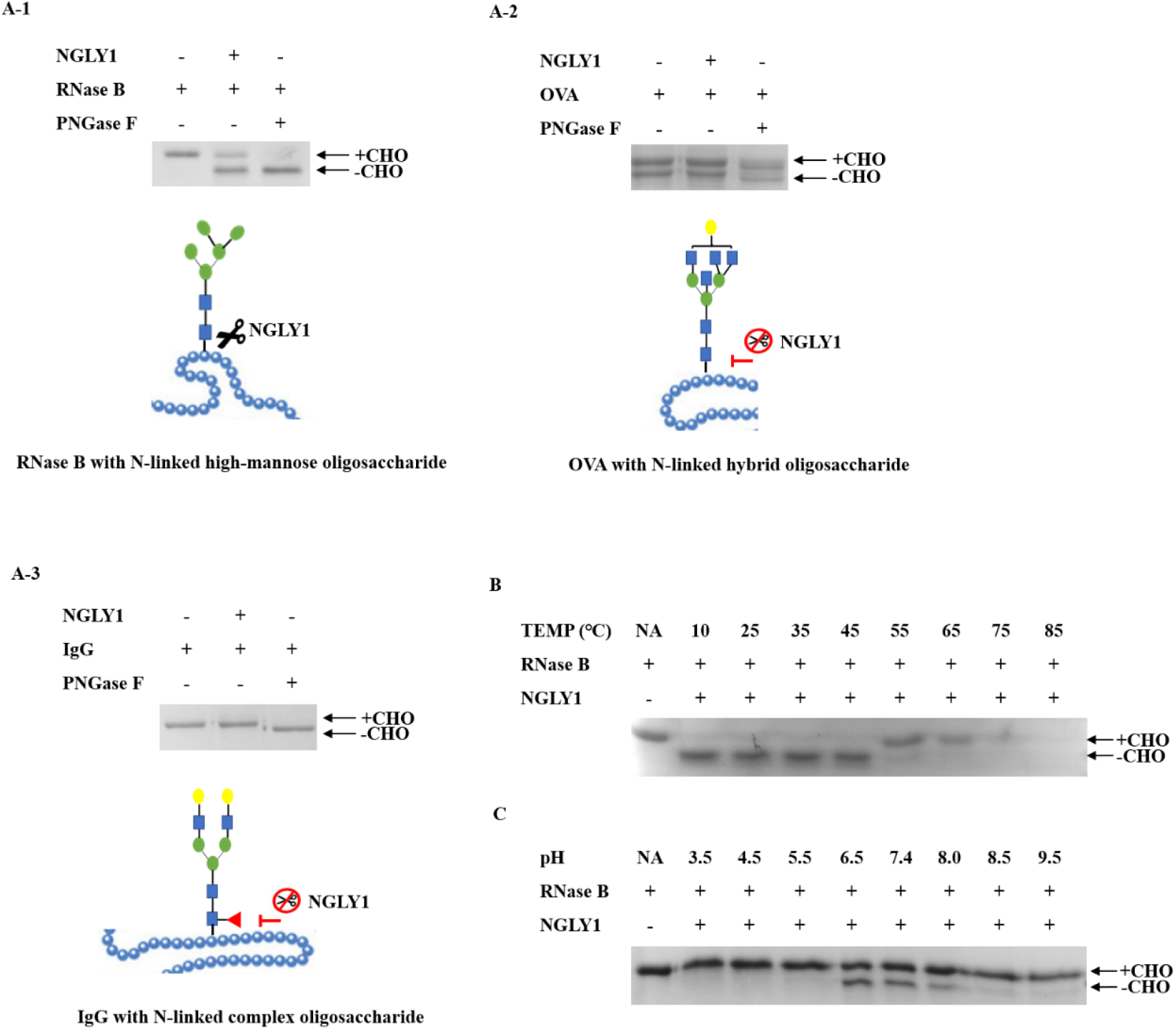
The effect of reaction conditions on NGLY1. (A-1) SDS-PAGE analysis of RNase B treated with NGLY1. (A-2) SDS-PAGE analysis of OVA treated with NGLY1. (A-3) SDS-PAGE analysis of IgG treated with NGLY1. The schematic diagram of RNase B, OVA, and IgG is shown (Aoyama et al., 2019; Sun et al., 2015; Wang et al., 2018a). Structure formulas: blue square, N-acetylglucosamine; green circle, mannose; yellow circle, galactose; red triangle, fucose. (B) The enzymatic reaction temperature of NGLY1. NGLY1 and RNase B were mixed and incubated for 12 h at different temperatures. (C) Enzymatic reaction pH of NGLY1. NGLY1 and RNase B were mixed, incubated for 12 h at different pH. NA: substrate control, without treatment; “-”: without NGLY1.

Three representative N-glycoproteins were digested with NGLY1 and PNGase F, including RNase B with N-linked high-mannose oligosaccharide, OVA with N-linked hybrid oligosaccharide and IgG with N-linked complex oligosaccharide. The deglycosylated forms of the substrate were detected by the downward shift of the glycoprotein band observed by SDS-PAGE analysis. In addition, the results revealed that NGLY1 could release N-linked glycans from RNase B (Fig. 2A-1) but could not catalyze the deglycosylation of OVA (Fig. 2A-2) or IgG (Fig. 2A-3). Based on these results, the optimum reaction conditions for NGLY1 were pH 7.4, temperature 37°C, and substrate RNase B.

### Enzymatic activities of NGLY1 functional active site R328 mutations

Our group reported a new NGLY1 mutation, R328G, which caused loss of N-glycanase function (Abuduxikuer et al., 2020). The established assay was used to determine the enzymatic activity of the NGLY1-R328 mutations qualitatively. A schematic diagram of the construction of 19 types of R328 NGLY1 mutations was presented in Fig. 3A. The results showed that 11 mutations were dysfunctional, including R328A, R328D, R328E, R328F, R328G, R328I, R328P, R328V, R328W, and R328Y, whereas NGLY1 mutants with R328H and R328T remained active as wild-type NGLY1. Seven mutations showed decreasing activity, including R328C, R328K, R328L, R328M, R328N, R328Q, and R328S (Fig. 3B). Quantitative analysis of enzyme activity for active mutations (R328H, R328K, R328C, R328L, R328M, R328N, R328Q, R328S, and R328T) was performed (Table 2). The enzyme digestion efficiency at 1 h was lower than that at 12 h. Therefore, we speculated that a 12 h reaction time should be used to determine the activity of NGLY1 mutations. Combined with WES or WGS, this in vitro enzymatic assay could provide a standard protocol for diagnosing NGLY1-CDDG.

**Table 2.**
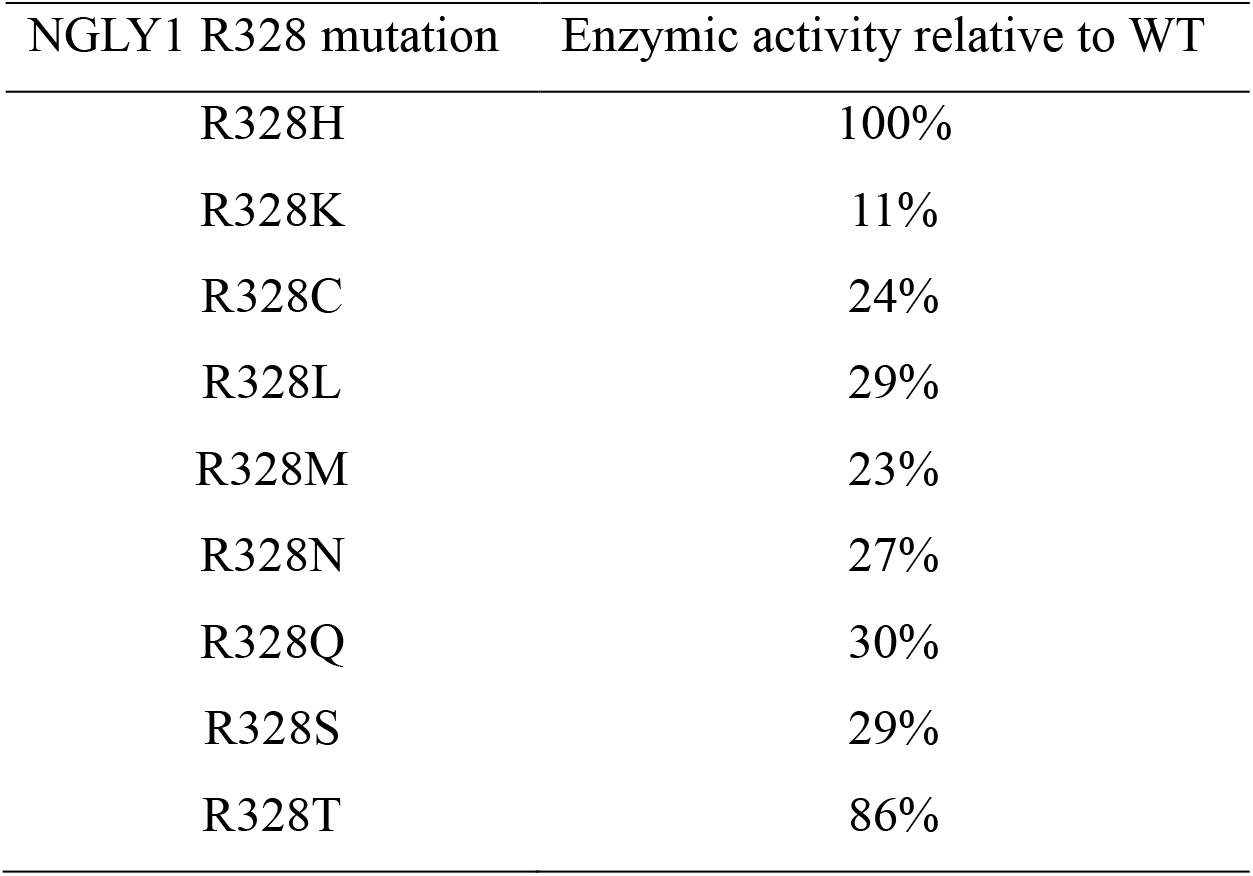
Relative activity of NGLY1-R328 mutants used in the study

| NGLY1 R328 mutation | Enzymic activity relative to WT |
| --- | --- |
| R328H | 100% |
| R328K | 11% |
| R328C | 24% |
| R328L | 29% |
| R328M | 23% |
| R328N | 27% |
| R328Q | 30% |
| R328S | 29% |
| R328T | 86% |

**Figure 3.**
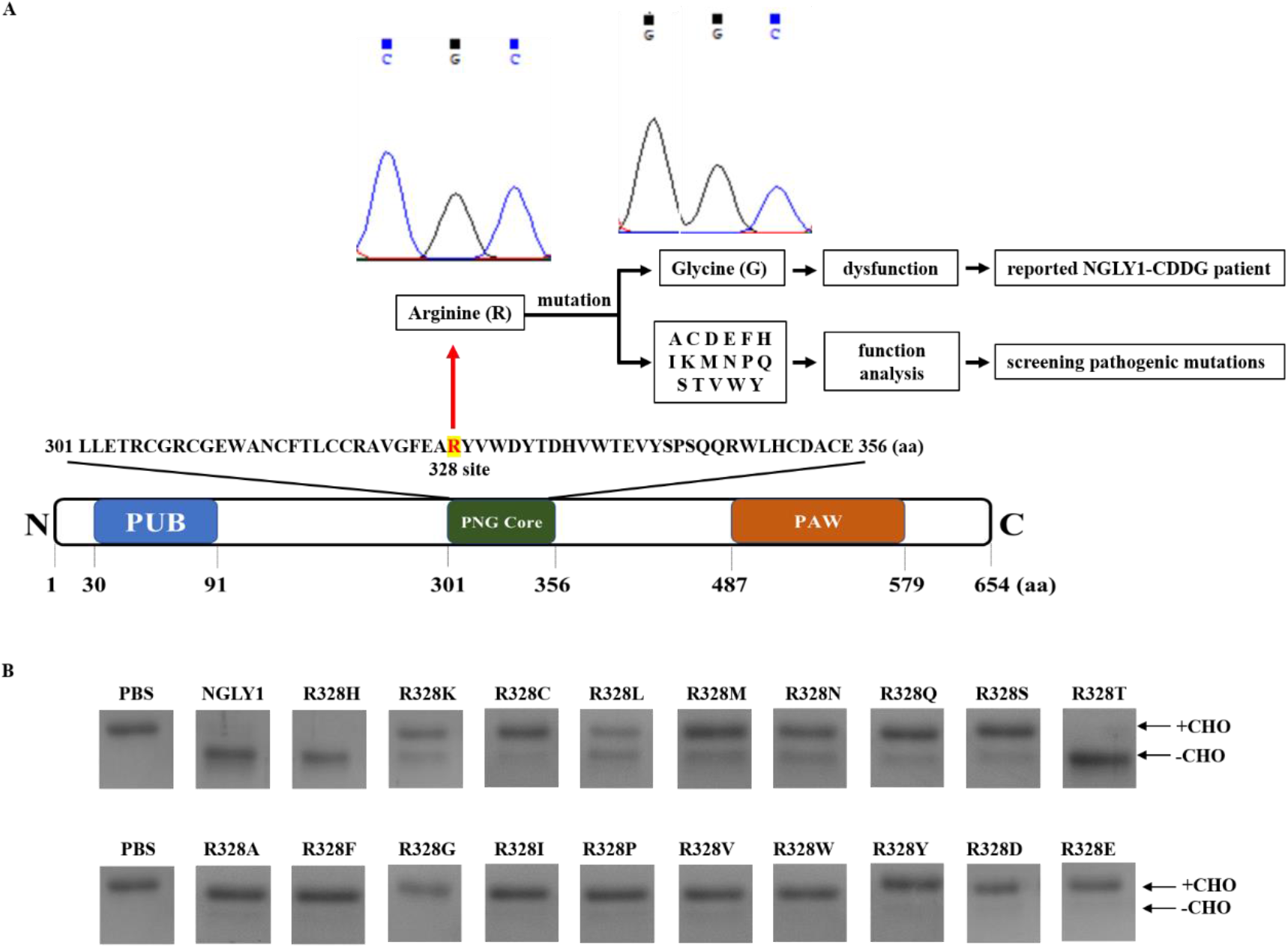
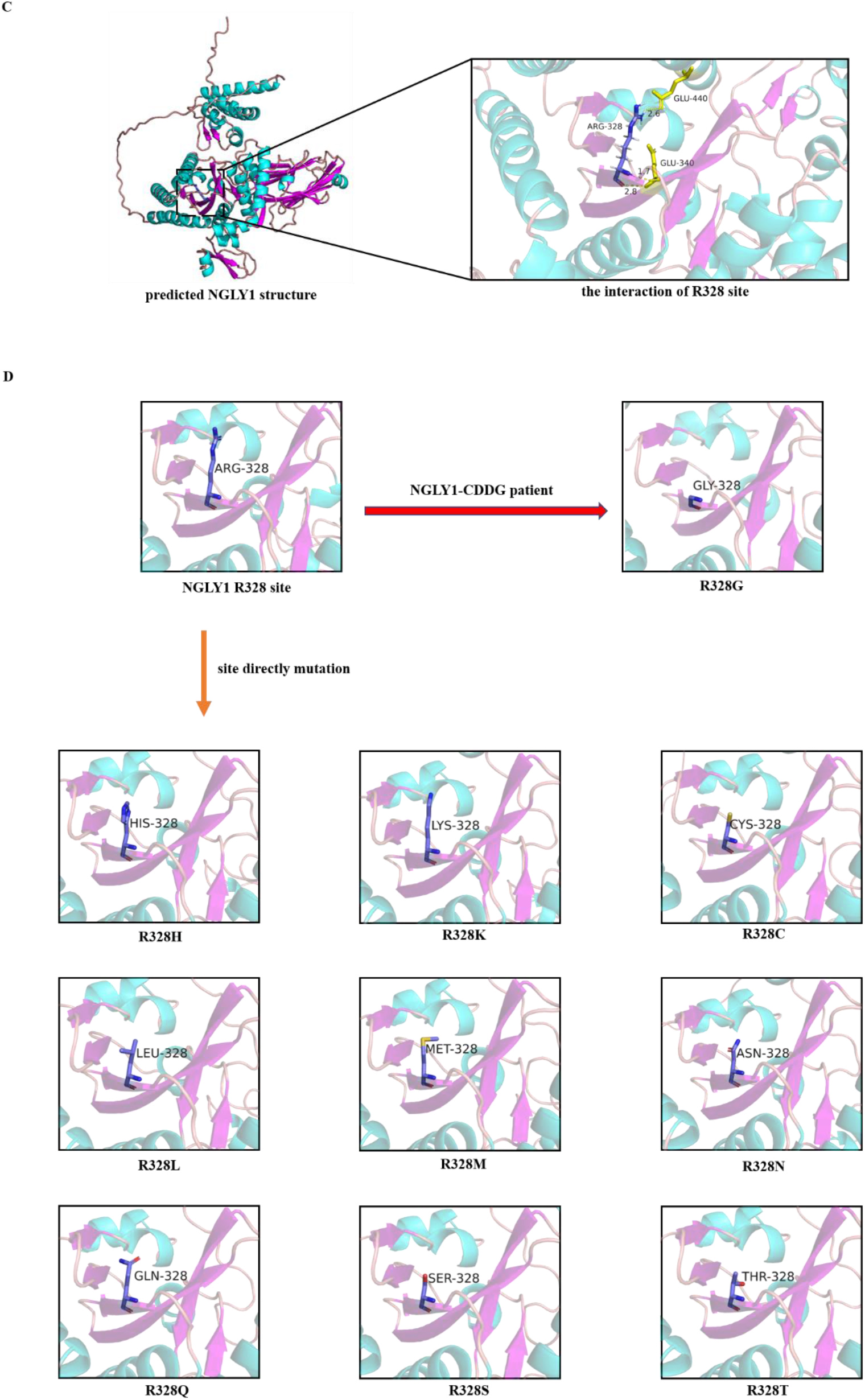
SDS-PAGE analysis of RNase B treated with NGLY1-R328 mutations, and the simulated structures of wild-type and mutated NGLY1 predicted by AlphaFold2. (A) The construction of the R328 site in NGLY1. (B) The enzymatic activity of mutations on the R328 site of NGLY1. (C) The interaction of the R328 site to other amino acids (E340 and E440). (D) The predicted structures of mutated NGLY1 (with enzymatic activity).A: alanine; C: cysteine; D: aspartic acid; E: glutamic acid; F: phenylalanine; G: glycine; H: histidine; I: isoleucine; K: lysine; L: leucine; M: methionine; N: asparagine; P: proline; Q: glutamine; R: arginine; S: serine; T: threonine; V: valine; W: tryptophan; Y: tyrosine.

We also analyzed the 3D structure of wild-type and mutant NGLY1. The simulated 3D structure of NGLY1 obtained using Alphafold2 showed three domains, among which the structure of the PUB domain was consistent with that revealed in a previous report (PDB ID: 2CM0) (Allen et al., 2006) (Fig. 3C). The simulated structures of mutated NGLY1 (possessing enzymatic activity) predicted by AlphaFold2 were presented in Fig. 3C and 3D, and the structures of dysfunctional mutations were also predicted (data not shown). Based on the predicted structure, the R328 site was critical for packing the catalytic domain (PNG Core domain of NGLY1). The R328 site interacted with E340 and E440 (upper part of Fig. 3C), which were conserved residues in NGLY1 (*Homo sapiens*), Ngly1 (*Mus musculus*), and YPNG1 (*Saccharomyces cerevisiae*). Disruption of such packing, either at sterically very small (R328A, R328G) or large residues (R328F, R328W) or electrostatically (negatively charged, R328D, R328E), would affect the catalysis. It was speculated that the structure of the amino acid side chain could influence the function of NGLY1.

### Effect of chemical compounds on NGLY1 activity

Since there is no effective treatment for NGLY1-CDDG, the compounds for promoting NGLY1 activity might be detected based on the established in vitro enzymatic assay. In addition, 402delR (Enns et al., 2014) and R328L mutations found in an NGLY1-CDDG patient were also evaluated. Several chemical compounds were added to the reaction system, including a reducing agent (NAC and DTT), an enzyme inhibitor (Z-VAD-FMK), and a denaturant (SDS) (Table 3). The results were analyzed by SDS-PAGE (Fig. 4). At concentrations of 0.1, 0.5, and 1 mM, NAC promoted the NGLY1 enzymatic activity (Fig. 4A-1), while NAC had no significant effect on NGLY1 with 402delR (Fig. 4A-2) and R328L (Fig. 4A-3). However, enzymatic activity of NGLY1, 402delR- and R328L-carrying NGLY1 was inhibited when NAC was added at a concentration of 5 mM (Fig. 4A). NGLY1 activity was increased by DTT (Fig. 4B-1), while the activity of the NGLY1 mutant harboring 402delR (Fig. 4B-2) and R328L (Fig. 4B-3) was inhibited. The enzymatic activity of NGLY1 was inhibited by Z-VAD-FMK (Fig. 4D-1) and SDS (Fig. 4D-2). In general, Z-VAD-FMK inhibits pan-caspase and yeast PNGase (Pagano et al., 2020).

**Table 3.** Effect of chemical compounds on wild-type NGLY1

| Chemical compounds | Activity | Metal ions | Activity |
| --- | --- | --- | --- |
| NAC | ↑ | BaCl<sub>2</sub> | ↑ |
| DTT | ↑ | CaCl<sub>2</sub> | ↑ |
| Z-VAD-FMK | ↓ | MgCl<sub>2</sub> | ↑ |
| SDS | ↓ | ZnCl<sub>2</sub> | ↓ |

↑: increase in enzymatic activity of NGLY1; ↓: decrease in enzymatic activity of NGLY1.

**Figure 4.**
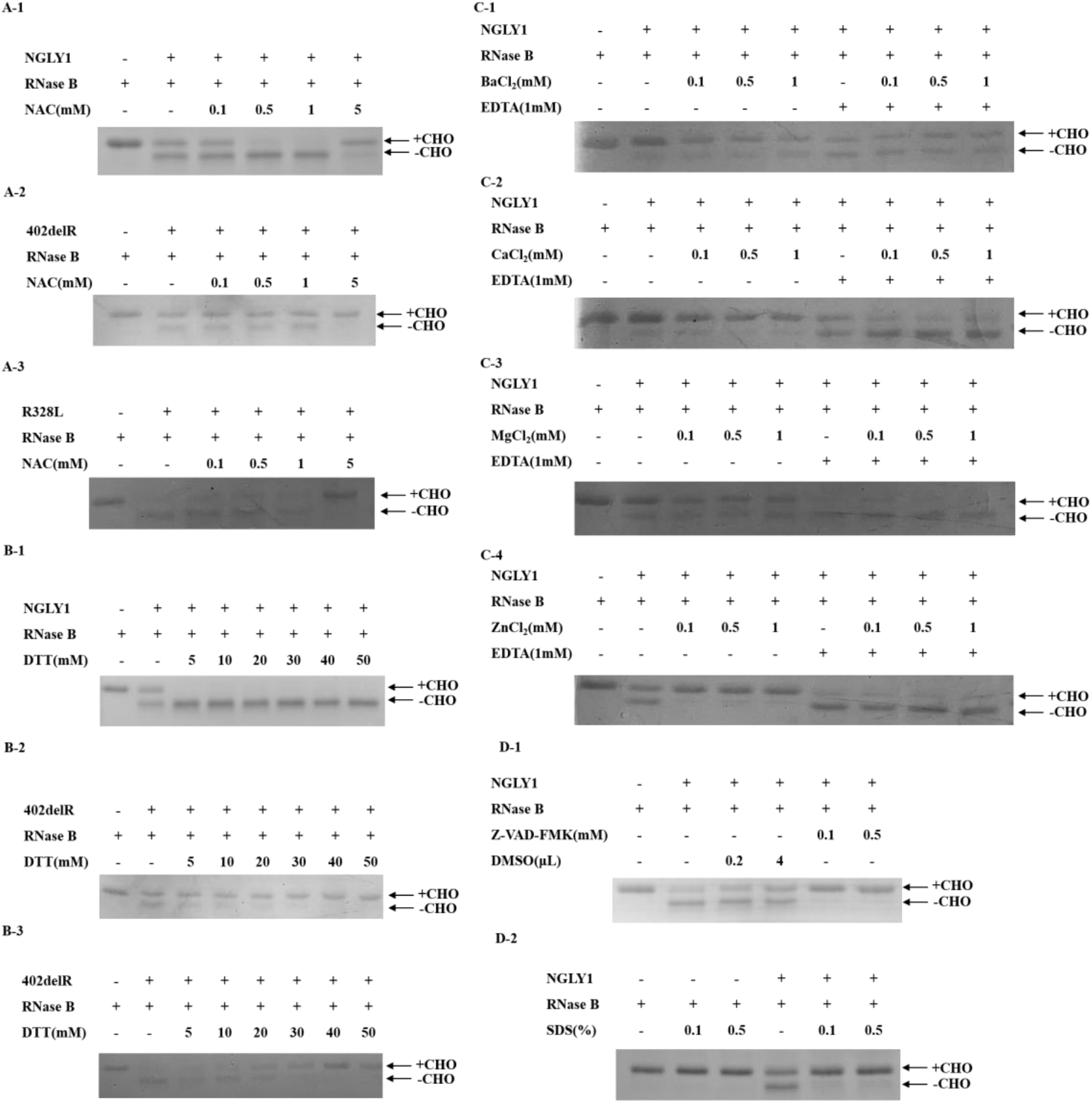
The enzymatic activity of NGLY1 treated with different chemical compounds. (A-1) NGLY1 treated with NAC. (A-2) NGLY1 mutant with 402delR (NGLY1-CDDG mutation) treated with NAC. (A-3) NGLY1 mutant with R328L (NGLY1-CDDG mutation) treated with NAC. (B-1) NGLY1 treated with DTT. (B-2) NGLY1 mutant with 402delR (NGLY1-CDDG mutation) treated with DTT. (B-3) NGLY1 mutant with R328L (NGLY1-CDDG mutation) treated with DTT. (C-1) NGLY1 treated with BaCl_2_. (C-2) NGLY1 treated with CaCl_2_. (C-3) NGLY1 treated with MgCl_2_. (C-4) NGLY1 treated with ZnCl_2_. (D-1) NGLY1 treated with Z-VAD-FMK. (D-2) NGLY1 treated with SDS.

Metal ions were also added to the enzymatic reaction (Table 3). The effects of metal ions on NGLY1 enzymatic activity were different (Fig. 4C). The activity of NGLY1 in digesting RNase B was promoted by BaCl_2_, CaCl_2_, and MgCl_2_ (Fig. 4C-1, 4C-2, and 4C-3). NGLY1 activity was inhibited by ZnCl_2_ (Fig. 4C-4); this result is consistent with previous reports of PNGase in mice (Suzuki et al., 1994) and *Caenorhabditis elegans* (Kato et al., 2007).

## Discussion

NGLY1-CDDG, the first reported congenital disorder of deglycosylation, is an extremely rare autosomal recessive disorder caused by pathogenic mutations in the *NGLY1* gene (Rios-Flores et al., 2020). NGLY1-CDDG patients are routinely diagnosed using WES or WGS (Haijes et al., 2019; Lipinski et al., 2020; Need et al., 2012; Rios-Flores et al., 2020). However, in the “HUABIAO” whole-exome public database of unrelated healthy samples, published by Hao et al. (Hao et al., 2021), 102 mutations of *NGLY1* are summarized (https://www.biosino.org/wepd/search?queryWord=NGLY1), of which 64 mutations are located in the exon of *NGLY1*, including 17 non-amino acid change mutations.

The *NGLY1* mutations detected by WES or WGS were not all related to NGLY1-CDDG. Therefore, a standard assay for the correlation between gene mutations and NGLY1 enzymatic activity is necessary to screen and diagnose NGLY1-CDDG.

In this study, an in vitro enzymatic assay system was established to evaluate the enzymatic activity of NGLY1 using MALDI-TOF MS, CE, and SDS-PAGE. NGLY1 and PNGase F can cleave intact N-glycans from N-glycoproteins (Sun et al., 2015). In addition, the CE atlas proved that NGLY1 and PNGase F could hydrolyze N-glycans from healthy human serum samples.

The highest mutation frequency of NGLY1-CDDG patients was presented in the nonsense mutation c.1201A>T (p.R401X), while only three-type mutations occurred at the R328 site in the PNG Core domain of NGLY1, including R328G (Abuduxikuer et al., 2020), R328C (Dabaj et al., 2021), and R328L (unreported patient); this result suggested that the R328 site was the core active site of NGLY1. Full-spectrum mutation libraries of the R328 site mutations on NGLY1 were constructed to demonstrate the relationship between the genotype and phenotype of NGLY1-R328 mutations. When R328 was mutated to an amino acid with the same charge (electropositivity), the mutated NGLY1 remained active, whereas the mutated NGLY1 (electronegativity) was dysfunctional. For mutations with electroneutrality, the structure of the amino acid side chain influences the function of NGLY1. Therefore, it was speculated that the electrification and side chain of amino acids possibly influenced the function of NGLY1. Some studies have indicated that electrification (Noda et al., 2020) and side-chain structure (Pappa et al., 2012) of amino acids would affect the function of proteins. Among the 19 mutations obtained, only 5 resulted from a one-nucleobase change (including R328C, R328G, R328H, R328L, and R328P) and were more likely to occur than other mutations. However, it was difficult to perform an experiment on quantitative enzyme activity. In this study, we preliminarily quantitatively analyzed enzyme activity through the gray value of two bands (RNase B + CHO and RNase B - CHO); however, this method was not accurate enough. The constructed full-spectrum functional mutation libraries provided a new diagnostic method for proteins mutated at R328 and other sites in NGLY1-CDDG and provided new insights into other single-gene disorders. Subsequently, we will analyze the correlation between the reported mutations in NGLY1-CDDG patients and NGLY1 enzymatic activity and the effect of R328 mutations on cells.

Several reported NGLY1 mutations in NGLY1-CDDG patients were analyzed using the established in vitro enzymatic assay system (including W244R, L318P, 402delR, and R469X), and the degrees of loss of mutated NGLY1 varied (data not shown). In this study, the effects of several compounds on NGLY1 enzymatic activity were analyzed using an established in vitro enzymatic assay. This in vitro assay may serve as a practical screening strategy for drugs and compounds with potential therapeutic value for NGLY1-CDDG patients. NAC and DTT have been shown to promote the enzymatic activity of wild-type NGLY1. NAC has been used as a mucolytic agent for treating airway muco-obstructive diseases (Ehre et al., 2019), and it might be a potential treatment for psychiatric disorders, including Alzheimer’s dementia, drug-induced neuropathy, and progressive myoclonic epilepsy (Minarini et al., 2017). In addition, the effect of DTT has been proven to mitigate hepatic and renal injury in bile duct-ligated mice and ameliorate hematopoietic and intestinal injury in irradiated mice (Li et al., 2019). Interestingly, the effects of NAC and DTT were not consistent among NGLY1-CDDG patient mutations. This finding is a cautionary warning of the need for precision medicine, in which treatment varies between patients with different NGLY1 mutations. Based on the established in vitro enzymatic assay system, drug screening and new potential drugs could be developed for NGLY1-CDDG therapy.

The diagnostic efficiency of NGLY1-CDDG could be improved through the combination of gene sequencing and an in vitro enzymatic assay system. However, further studies are required to determine the correlation between the degree of NGLY1 inactivation and the clinical phenotype of NGLY1-CDDG patients.

